# The chromosomal basis of species initiation: *Prdm9* as an anti-speciation gene

**DOI:** 10.1101/170860

**Authors:** Donald R. Forsdyke

## Abstract

Mechanisms initiating a branching process that can lead to new species are broadly classified as chromosomal and genic. Chromosomal mechanisms are supported by breeding studies involving exchanges of individual chromosomes or their segments between mouse subspecies. There are also studies of the rapidly mutating mouse *PR/SET-domain 9* (*prdm9*) gene, which encodes PRDM9, a protein targeting DNA recombination hotspots. When PRDM9 is bound symmetrically with equal strength, the meiotic repair of mutations in one parental strand, based on information on the allelic strand (conversion), would seem to be unbiased in discriminating between strands. So mismatches detected between pairing paternal and maternal DNA strands (heteroduplexes) undergo unbiased conversions (to homoduplexes). This leaves uncertainty on whether a mutation has been corrected or compounded. However, it has been hypothesized that tagging of mismatch regions, so that both strands are epigenetically marked as uncertain, would make it possible over numerous generations for mutations to be corrected (biased conversions) whenever *asymmetry* is detected. Thus, variation would decrease and members of a species would remain within its bounds. Intriguingly, new experimental studies show that, when chromosomally interpreted, PRDM9 also works through *asymmetrical* epigenetic labelling to confine members to species bounds. To the extent that the experimentally observed and hypothetical anti-speciation asymmetries can be related, chromosomal mechanisms are further supported.

## Introduction

Hypotheses on the initiation of a branching process that can lead to new species are broadly classified as chromosomal and genic [1–4]. Agreement is sought as to which initiation mechanisms are actually, rather than hypothetically, capable of originating species, and which are most likely to have operated in the general case [5–7]. Chromosomal hypotheses invoke sequence disparities between parental chromosomes so that meiotic pairing of ‘homologous’ chromosomes fails within the gonad of their offspring (hybrid). Thus, there can be no exchange of DNA segments (recombination) and the production of gametes ceases (hybrid sterility). Arising from studies of the *PR/SET-domain 9* gene (*prdm9*), there is now “fresh evidence for a genetic connection between recombination and hybrid sterility,” suggesting “the intriguing possibility that recombination and speciation are mechanistically coupled” [8]. However, the ‘speciation genes’ commonly invoked to explain this have become increasingly elusive.

Specific genes affecting species initiation have been proposed for fruit fly [9]. While the existence of similar genes in other species is doubted [10–12], the *prdm9* gene that is expressed in early germ cell maturation [13, 14] is now thought a likely “first mammalian candidate for a speciation gene” [15]. More definitely, it is “the only mammalian speciation gene yet identified” [16]. However, studies to be reviewed here suggest that its major role is to encode a genome maintenance protein that works to *retain a line of organisms within species bounds*. Thus, it is better regarded as an *inhibitor* of speciation – an anti-speciation gene – that opposes speciation when initiated chromosomally [17, 18]. Whatever the outcome, “cracking the curious case of PRDM9 promises to provide important insights into large swaths of biology, from human genetics to speciation” [19].

This review has three parts. The first describes the role of the *prdm9* protein product, PRDM9, in adding epigenetic methylation marks to histones. This designates the chromosomal location of meiotic recombination hotspots where there can be an *asymmetric* conversion of information in one parental genome to that of the other. However, like some other genes whose products target DNA, *prdm9* is on a mutational ‘treadmill,’ being forced to change rapidly to keep pace with DNA sequence changes. Its DNA target ‘calls the tune,’ yet in a paradoxical way. Hotspots appear self-destructive in that a given PRDM9 protein faces, generation after generation, diminishing target availability, thus ‘obliging’ its gene to rapidly mutate in response to a so far unexplained evolutionary pressure for the designation of new targets. The second part deals with the new light cast on chromosomal speciation by experimental modifications of PRDM9 that confers on it the ability to tune-call, so shifting hotspot locations. Finally, the possibility is considered that these observations relate to a postulated asymmetry requirement of the epigenetic marking of sequences whose accuracy is in doubt [18].

## PRDM9 and recombination hotspots

### Histone modifications by PRDM9

Of the barriers with a potential to initiate the reproductive isolation needed for sympatric divergence of one species into two, the hybrid sterility barrier resulting from defective meiosis can *only* play a primary role. Any barrier can be primary, but there is an onus upon those believing another barrier (e,g, hybrid inviability) to be primary, to show that it had not been preceded by a period, however brief, of hybrid sterility [20]. As for mechanisms, although controversial there is now growing evidence that nucleic acid “pair-first” (rather than “break first”) models accord well with meiotic recombination as described in many organisms [21–25]. Subversion of such pairings by sequence disparity would create conditions favoring speciation.

Furthermore, while cellular nucleic acids are closely associated with proteins that could affect recombination-related nucleic acid interactions, many aspects of DNA pair-first mechanisms can be reproduced *in vitro* in the absence of proteins [26, 27]. Assuming nucleic acid evolution to have preceded that of proteins [28, 29], one can envisage the kinetics of recombination – dependent on nucleic acid chemistry [30] – to have been affected by later evolving proteins.

Thus, in practice, pair-first mechanisms cannot really be first. There is need for a prior modification of proteins, such as the histones around which DNA is organized as nucleosome complexes. Modification of such nucleoprotein (chromatin) complexes to improve access to DNA is one of the functions of PRDM9, the zinc finger part of which recognizes with high specificity a short DNA sequence motif at the nucleosome surface, so defining a 1-2 kb recombination ‘hotspot’ region. This recognition activates the protein’s PR/SET domain that catalyzes the local addition of methyl groups to certain lysine residues of one of the histone proteins (H3). Trimethylation of H3 generates a ‘nucleosome-depleted region’ where recombination is facilitated (i.e. homologous strands can pair, there is branch migration to increase the area scanned, and the enzyme causing double-strand breaks, SpoII, can be recruited).

### Conversion restrains diversity

That meiosis, apart from generating diversity by shuffling parental genomes, also has the potential to *restrain* diversity, has a long history. In 1901 Montgomery proposed that meiosis would “rejuvenate” the colorfully staining material – chromatin – in disparate chromosomes [31]:

When the spermatozoon conjugates with the ovum there is a mixture of cytoplasm with cytoplasm, of karyolymph with karyolymph, … but there is no intermixture of chromatin, for the chromosomes … remain more separated from one another than at any other stage …. But after this beginning stage of the germinal cycle, … in the synapsis stage … we find a positive attraction between paternal and maternal chromosomes. The reason for the final union of these chromosomes is obvious; it is evidently to produce a rejuvenation of the chromosomes. From this standpoint the conjugation of the chromosomes in the synapsis stage may be considered the final step in the process of conjugation of the germ cells. It is a process that effects the rejuvenation of the chromosomes; such rejuvenation could not be produced unless chromosome of different parentage joined together, and there would be no apparent reason for chromosomes of like parentage to unite.

That the rejuvenation could involve exchange of linear information between disparate chromosomes was outlined by Winge in 1917 [32], and later elaborated in terms of DNA mismatch repair [1, 30, 33-34]. If we assume rejuvenation to imply a restoration (“correction”) of DNA sequences that have strayed from the species norm (i.e. they are to be returned closer to their formerly ‘pristine’ base composition and order), then it follows that a *lack* of such rejuvenation would allow DNA differences to accumulate, so possibly leading to reproductive isolation and speciation. Rejuvenation would restrain diversification, and hence, speciation. Fundamental to this are mechanisms of meiotic recombination.

Meiosis within a parental gonad generally provides for the unbiased distribution of grandpaternal and grandmaternal genomic information among offspring. Sometimes detected as a failure of characters to follow a Mendelian pattern of distribution, the *asymmetrical* conversion of information from one genome into that of the other has been called “gene conversion.” But, in principle, the asymmetry would also apply to non-genic sequences. Given two formerly homologous segments, A and B, which have come to differ slightly (i.e. are now heterologous), information in A can be converted into that of B, or vice-versa. Thus, regarding individual offspring, diversity is diminished. However, the direction of the conversion may be *generally* unbiased in that in one individual A may have converted to B, and in another B may have converted to A. Thus, at the population level, in the absence of conversion bias there will be no change in the frequencies of A and B. Diversity is not diminished.

On the other hand – rather than let natural selection be the arbiter of frequencies should one segment prove more advantageous than the other – *if there were sufficient information*, a parent could ‘place an educated bet’ on the conversion direction to adopt. For example, if it were ‘known’ that A was the species norm and/or that B had experienced a recent mutation, then there could be an appropriate conversion bias. Since most mutations are detrimental, resulting in negative selection in the line of organisms within which they occur, the best bet would be to convert B into A. If all parents were to bet similarly then diversity within the population would be diminished. Intriguingly, various studies show that a sequence motif recognized by PRDM9 (so defining a recombination hotspot) is directionally converted in mice heterozygous for that hotspot in favor of the sequence *less strongly* recognized by that protein (meiotic drive). I consider later the possibility of ‘educated bets’ in the light of studies of incipient speciation in mice.

### DNA inconstancy drives protein treadmill

Proteins often act as enzymes catalyzing changes in other molecules – their specific substrates – which may be micromolecules (e.g. glucose), or macromolecules (e.g. DNA). In the course of evolution there are changes in proteins as a consequence of mutations in the genes that encode those proteins. Their substrates do not usually participate in this directly. For example, as organisms evolve, specific enzymes may change and so improve the utilization of glucose. That’s natural selection. *But glucose itself has no say in this*. It does not change to lighten, or impede, the task of the enzymes. For some enzymes, however, DNA is a specific substrate. DNA both encodes the enzymes (locally in the regions of their genes), and is either their local or general substrate (target). However, DNA, by virtue of base changes, can change its character. Thus, it has the power to lighten or impede the tasks of the enzymes that act on it. More than this, the *constancy* of glucose means that the enzymes of glucose metabolism can count on glucose being the *same* from generation to generation. The *limited constancy* of DNA means that some of the proteins that react with it are on an evolutionary treadmill. They are “red queens” that must change as their substrate changes [35].

While some proteins target nucleic acids generically by virtue of their unchanging phosphate-ribose structures, others are not indifferent to the composition or order of bases. So, as nucleic acid bases change, the proteins that target nucleic acids more specifically may need to adapt functionally to ensure optimal binding to their substrate. This *adaptive treadmill* implies that, as lines diverge, the mutation rate in nucleic acid recognition proteins must be greater than in most other proteins [36]. If its genes encoding nucleic acid recognition proteins do not rapidly mutate, an organism may not survive. While here the DNA ‘dog’ may wag the protein ‘tail,’ sometimes the converse may apply. Changes in proteins encoded by certain “mutator” genes can exert a genome-wide effect on base composition [37]. Likewise, a mutated *prdm9* gene can, in one fell swoop, exert a genome-wide effect on the nature and location of target sites that focus recombination events (see below). Paradoxically, however, over evolutionary time it is usually these target sites that ‘call the tune’ resulting in the positive selection of *prdm9* mutants: “PRDM9 has been under selective pressure to switch to new targets. However, the reasons for this selective pressure remain mysterious” [35].

### The hotspot conversion paradox

The genomic locations of recombination events are usually not randomly distributed. In microorganisms there are stand-specific, mainly gene-located, “crossover hotspot instigator” (Chi) sequences that are recognized by enzymes with a role in recombination [38]. Multicellular organisms also have sequence hotspots where recombination most frequently initiates. Such hotspots may relate to genic boundaries (e.g. transcription start sites), either as a primary feature [39], or as a default option when sites (motifs) recognized by PRDM9 are not operative [40]. In organisms possessing a functional *prdm9* gene (e.g. primates, rodents) the primary hotspot option is often extra-genic or within introns. These DNA regions have more potential than exons to extrude kissing-loop structures [21, 30].

However, there is a conversion bias in favor of the *less strongly recognized* mutant parent genome. While, through the generations, members of the majority population continue to produce unmutated target sites (scheme at left), recurrent crosses with members of the growing mutant population (the cycle on the right; thick lines), erode this lead. Thus, the *prdm9* gene, in its X-recognizing form, becomes progressively irrelevant.

Chi being a conserved sequence in bacteria, it was at first thought that there would be a conserved, specifically targeted, binding sequence for PRDM9. Much attention was paid to a degenerate 13-base consensus motif that was often found in human hotspots. However, although recombination initiation hotspots may endure for many generations, eventually they disappear (erode) due to an unexplained *directional* conversion bias that accompanies recombination (Fig. 1). Thus, Wahls and Davidson [41] observed that “hotspots seed their own destruction,” and hence, to maintain recombination, fresh sites with different sequence characteristics must emerge. Focusing on yeast, which do not have the *prdm9* gene, they noted that “within the genome is a collection of hotspot-active DNA sites and a reservoir of ‘cryptic’ DNA sequence motifs that can be rendered active by as little as a single-base-pair substitution.” Under this “equilibrium dynamic model” the substitution refers to a DNA target motif, not to the enzyme that recognizes that motif. However, the possibility was entertained that for organisms with the *prdm9* gene, novel motifs could be so designated by its PRDM9 product if the gene were appropriately mutated (“*prdm9* shift model”). Accordingly, the widely distributed hotspot sequences that had eroded could be viewed as having ‘called the tune,’ and the gene had ‘responded’ by mutation (i.e. the mutation conferred some selective advantage). The latter view is now supported by extensive studies, such as those on the infertility of the male offspring of crosses between mouse subspecies carried out in the Forejt laboratory (see later).

**Figure. 1.**
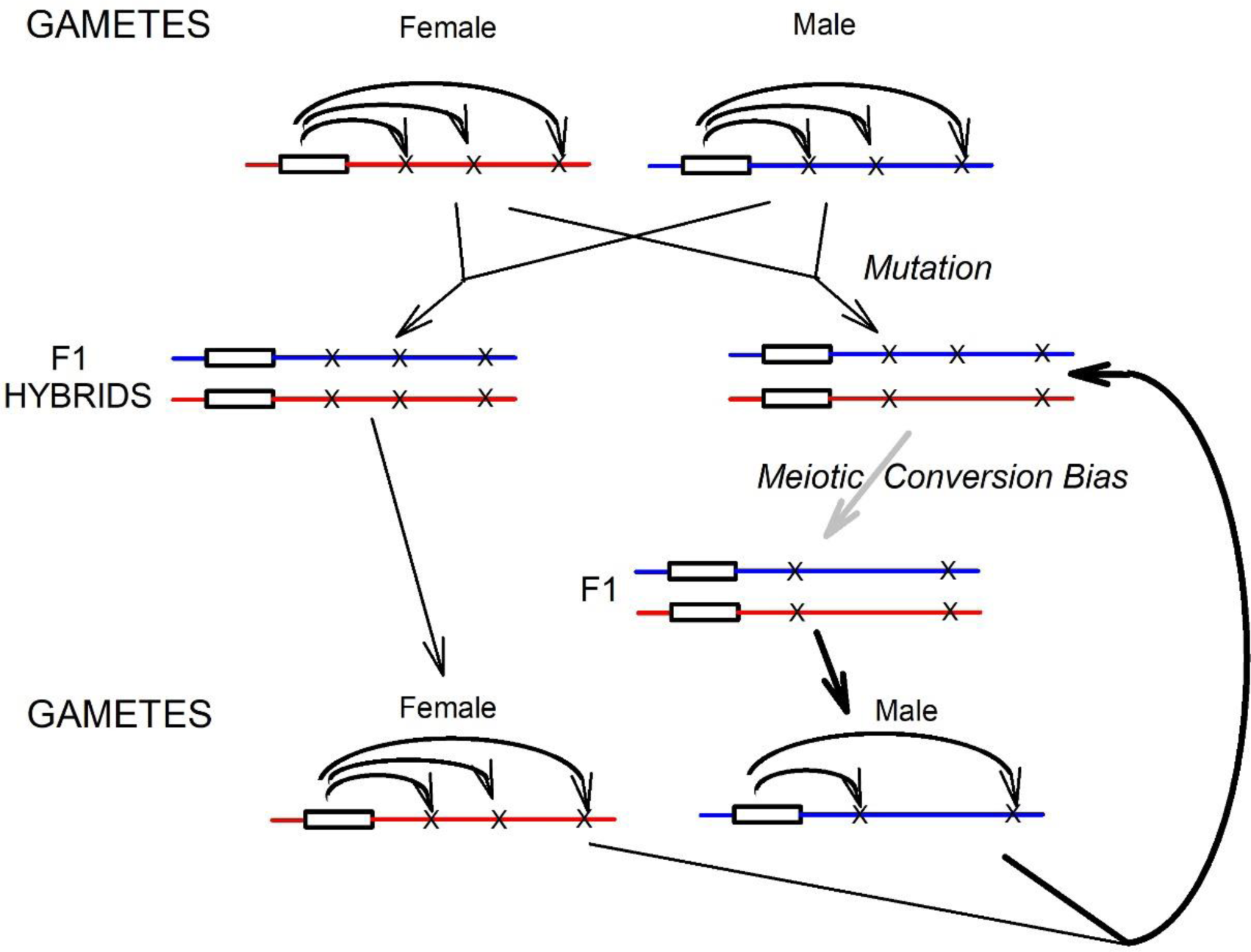
Cyclic erosion of PRDM9 binding sites during repeated within-species crosses. Homologous *prdm9* genes are shown as white rectangles flanked by horizontal lines (genomes; red and blue in the online version). At top, curved arrows indicate potential PRDM9 targets (X) at various genomic locations. At left, parental gametes unite to form normal F1 hybrids with symmetrical target sites. These hybrids, in turn, form parental-type gametes (the choice of sex is here arbitrary). At right, a mutation has changed a target site in one parental gamete (loss of X). Despite the local asymmetry, both normal and mutant sites are recognized at meiosis by PRDM9, and chromosomal pairing and recombination occur, albeit perhaps less efficiently.

When comparing diverging sequences, it is argued that that which is conserved between the sequences is evolutionarily important, whereas that which is less conserved is less important – indeed, that is why the ‘hand of evolution’ may have ‘chosen’ to discard it. Thus, when confronting the “hotspot conversion paradox” (Fig. 1), Boulton *et al.* noted [42] that since “biased gene conversion is a typical consequence of recombination at hotspots,” then “the sites thought to initiate crossing over cannot be maintained by the benefits of the events they cause”. In other words, change *itself* seemed to have an adaptive value.

So there are circumstances under which non-conservation seems to have a *greater* adaptive value than conservation. Likewise, proteins such as PRDM9 are themselves transient, displaying extensive variation over time. Their genes are on a treadmill, having to mutate to keep up with the constantly changing hotspot landscape. Thus, there is high within-species variation (e.g. polymorphism), and even greater between-species (e.g. human-mouse) variation, in the DNA recognition regions (zinc fingers) of *prdm9* genes. Grey *et al.* note [43]:

The PRDM9 gene is well-conserved among metazoans, however the domain encoding the zinc finger array experiences an accelerated evolution in several lineages, including rodents and primates. This accelerated evolution is restricted to codons responsible for the DNA-binding specificity of PRDM9 zinc fingers, which appear to have been subjected to positive selection.

## PRDM9 and speciation

### Subspecies as incipient species

Breeding studies from the Forejt laboratory, involving exchanges of individual chromosomes or chromosome segments between members of two mouse subspecies (*Mus musculus musculus* (*Mmm*) and *Mus musculus domesticus* (*Mmd*)), support chromosomal, rather than genic, disparity as a basis for species initiation [44–46]. The subspecies had not been designated as “species” because the reproductive isolation that defines species was incomplete in that a degree of fertility was still evident among female offspring (Haldane’s rule). The subspecies were incipient species with a potential eventually to attain full species status [47]. Two important strains within the subspecies are PWD (from *Mmm*) and B6 (from *Mmd*). They are estimated to have diverged from a common mouse ancestor about 0.5 million years ago and, while their DNA sequences are closely identical, they differ in their *prdm9* genes and the multiple specific targets of the PRDM9 proteins encoded by those genes. The latter targets can be represented (see Fig. 2, top) as Xs (for PWD) and black dots (for B6). These symbols represent the target sequence motifs *as they now exist* and, largely because of *past* erosions, they are likely to *differ* from versions that were active at earlier times (closer to the time of the initial PWD-B6 divergence).

**Figure. 2.**
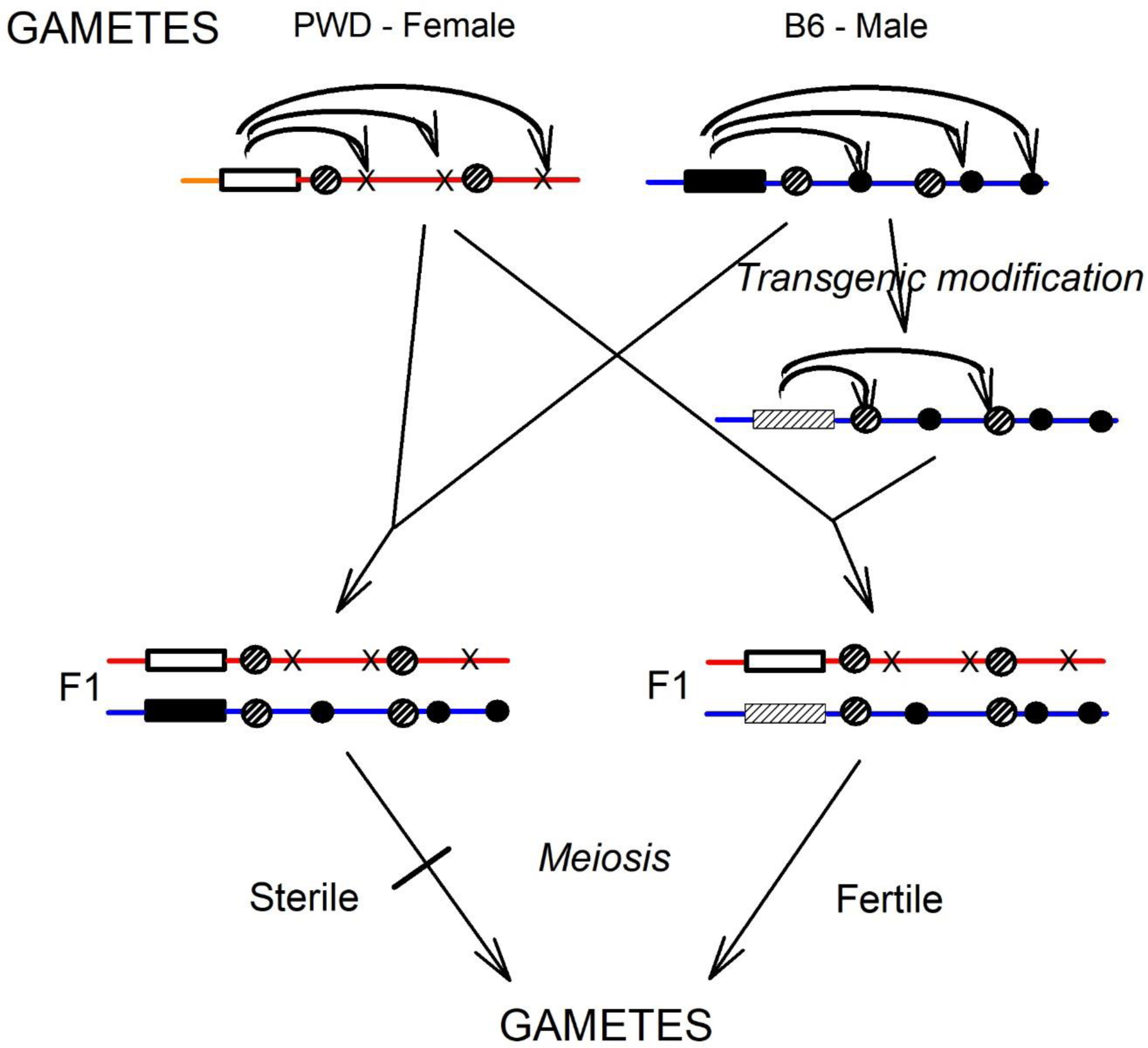
Hybrid sterility induced by pervasive reciprocal asymmetry of PRDM9 target sites when members of mouse subspecies are crossed (left), and its rescue in a transgene “humanized” strain (right). At top, subspecies-specific *prdm9* genes are shown as rectangles (PWD strain, white; B6 strain, black). Corresponding potential genomic targets (Xs and black dots) are indicated by curve arrows. At left, gametes unite to form F1 hybrids that develop and grow normally. However, due to a pervasive asymmetrical meiotic conversion bias that impairs gamete formation, they are sterile. At right, the B6 version of the *prdm9* gene has been transgenically modified (“humanized”) such that the zinc fingers no longer recognize B6 targets (black dots), but do recognize sequences elsewhere in the genome (striped dots) that happen to closely resemble the target of the human *prdm9* gene. When the humanized B6 mouse is crossed with a normal PWD mouse, meiosis proceeds normally due to the symmetry between targets (striped dots). This overcomes the effects of pervasive asymmetries related to the presence of the PWD version of *prdm9*, and the mice are fertile.

### Erosion of target motifs

A consequence of the erosion of specific target sites (Fig. 1), is that *today’s* PWD mice encode PRDM9 proteins with *less* affinity for their target sites on the PWD genome than for target sites at the corresponding positions on the B6 genome (the latter are not symbolised in Fig. 2, top right). In other words, *today’s* PWD-encoded PRDM9 proteins would, if given the opportunity, display *stronger* binding to a B6 genome than to a PWD genome. Likewise, *today’s* B6-encoded PRDM9 proteins would, if given the opportunity, display *stronger* binding to a PWD genome than to a B6 genome. That opportunity arises when members of the two subspecies are crossed.

For a *within*-subspecies cross (Fig. 1), the decreased affinity does not disturb the general *symmetry* of binding of PRDM9 proteins. So long as appreciable affinity is retained, there should be fertile offspring. However, when there is a *between*-subspecies cross (Fig. 2) the binding is generally *asymmetrical* and the story is very different. When a PWD female is crossed with a B6 male, all male offspring are sterile [48, 49]. A likely mechanism for this is shown at the left of Fig. 2. Because of the prior erosion, on the genome donated by the PWD gamete the target sites for the PRDM9 encoded by the PWD version of *prdm9* are weaker than the *same* target sites on the genome donated by the B6 gamete. Similarly, on the genome donated by the B6 gamete the target sites for the PRDM9 encoded by the B6 version of *prdm9* are weaker than the *same* target sites on the genome donated by the PWD gamete. The wide spread asymmetry is somehow sufficient to impede meiotic pairing and the individual is sterile.

### Humanized targets restore fertility

For within-subspecies crosses, binding site affinity progressively decreases over evolutionary time (Fig.1), with the potential to lead to sterility. Thus, individuals with *prdm9* mutations affecting the PRDM9 zinc fingers (so designating fresh binding sites), should have a selective advantage. Indeed, experimental exchanges of mouse *prdm9* genes indicate restoration of fertility [14]. An extreme version of this was to insert a human zinc-finger DNA recognition region into a mouse *prdm9* gene (Fig. 2, right). Given both the great length of their genomes, and the separation of humans and mice from their common ancestor some 150 million years ago, it was likely that there would be some symmetrical, non-eroded, potential target sequences in the mouse genome (indicated by striped dots in Fig. 2). These would be targeted, through its protein product, by the transgenically modified *prdm9* gene. B6 targets (black dots in Fig. 2, right) were denied a corresponding B6 *prdm9*-encoded protein, but the pervasive asymmetry of PWD targets should have been recognized by the corresponding PWD *prdm9*-encoded protein. The finding that the cross was fertile indicated a primary role for PRDM9 zinc-finger mutations in preserving within species fertility [16].

Thus, the eroding multiple target sites in a genome ‘call the tune.’ The *prdm9* gene must ‘respond’ to maintain fertility. This maintenance is in keeping with the observation of Oliver *et al.* for humans that “allelic variations at the DNA–binding positions of human PRDM9 zinc fingers show significant association with decreased risk of infertility” [50]. Davies *et al.* concluded that “the full fertility of humanized mice implies there are unlikely to be any specific essential PRDM9 binding sites” [16].

## Epigenetic marking of suspect sequences

### An anti-speciation gene

From generation to generation, growing DNA sequence diversity between two intraspecies groups makes speciation more likely [51]. Conversely, anything that facilitates the transgenerational uniformity of genomic information should make speciation less likely. As mentioned above, in many quarters *prdm9* is considered a ‘speciation gene,’ indicating its direct *positive* involvement in the speciation process, such as by *increasing* the sequence diversity that can lead to hybrid sterility. Indeed, PRDM9 can work to induce sterility as shown in Fig. 2 (left side). However, concerning this positive involvement, Davies *et al.* remark that “the rapid evolution of the zinc finger array of PRDM9 implies an unexpected transience of this direct role” [16]. In keeping with this, Fig. 2 (right side) raises the possibility that the development of sterility would be forestalled by zinc finger mutations (of which the humanized version is an extreme example) that would work to reduce variation and produce fertile offspring. The sterility could indeed be evolutionarily transient. Supporting this view, human PRDM9 zinc finger mutations have been found to associate with *protection* against hybrid sterility [50, 52]. Thus, PRDM9 can be viewed as normally promoting recombination, hence providing opportunities for repairing mutations and so impeding the sequence divergence that can lead to speciation. It opposes the hybrid sterility that is “characterized by failure of pairing (synapsis) of homologous chromosomes and an arrested meiotic prophase owing to lack of repair of recombination intermediates” [16]. In this light, *prdm9* is best regarded as an *anti-speciation* gene in that it facilitates the correction of meiotic mismatches (heterozygosities). There is now diminishing support for the view that adverse genic interactions (epistasis), referred to as Dobzhansky-Muller incompatibilities, are as fundamental to speciation as meiotic chromosomal mismatches [12, 53]. Interactions between fruit fly examples likely evolved late in the speciation process [9, 54].

### A tale of two asymmetries

Yet none of this easily explains the “hot spot paradox” [42], namely the erosion of PRDM9 target sites when asymmetrically disposed between parental genomes (Fig. 1). For some reason, the *prdm9* gene is on a treadmill being constantly ‘obliged,’ through mutation, to reinvent itself, so encoding a novel PRDM9 protein product that can summon up fresh, symmetrical, target sites. These sites then become epigenetically marked through histone methylation.

In another context, the epigenetic symmetry/asymmetry issue also emerged with a hypothetical mechanism for transgenerational error-correction through biased meiotic conversion [18, 27]. Here a means by which a repair process could achieve a gradual (multi-generational) bias against mutations was envisioned. When meiotic branch migration encounters a point mutation, a base pair mismatch arises and is repaired, so either the maternal or the paternal base is converted to the complement of its opposing base in the heteroduplex. In the absence of any reason to expect a systematic bias in this repair process, chance alone would seem to determine a mutation’s fate. However, it is supposed that where a meiotic strand exchange first encounters a base pair mismatch (indicating a likely mutation) *both* strands are epigenetically marked to designate uncertainty. Such a mark, if retained within the new organism’s germ cell line, will convey the message “there is a near 50% chance that this allele is a mutation.” If this epigenetic tag could promote subsequent meiotic strand exchange, and invite the next meiotic partner (less likely to harbor a mutation) to serve as the donor strand in the subsequent mismatch repair, this could ensure that over many generations biased conversions will remove mutations. Such a process, by reducing the rate of mutation-driven genetic divergence within breeding populations, would act as a force against speciation. Fig. 3 outlines the proposed tagging principle, which has some resemblance to the epigenetic tagging of damaged DNA in cell lines [55].

**Figure. 3.**
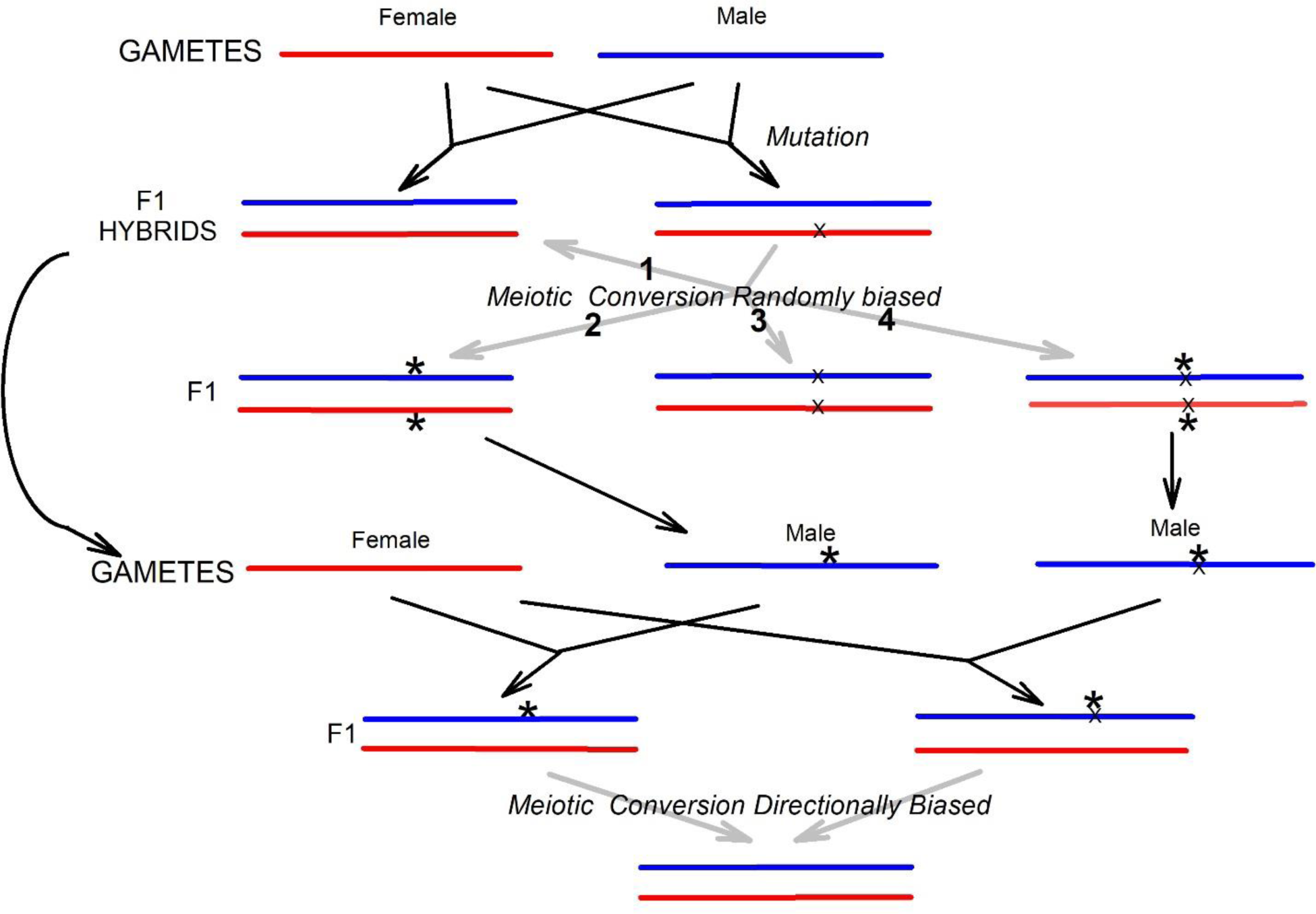
Marking as “suspect” a sequence whose accuracy is in doubt facilitates remedy (albeit delayed) by directional conversion bias. Resulting from a normal cross (top left), healthy homozygous F1 hybrids are representative of the species, members of which produce normal gametes (curved arrow). However, a mutation (X) in one parent (top right) may produce a heterozygous F1 hybrid. If the mutation is such that mutant and non-mutant strands cannot be distinguished, meiotic conversion bias will occur randomly, with four possible outcomes (1, 3, 2 and 4). 1: Bias in favor of the unmutated strand *corrects* the mutation. 3: Bias in favor of the mutated strand *compounds* the mutation. 2 and 4: In the region of the heterozygosity, conversion is accompanied by epigenetic tagging as “suspect” (asterisks). This affects both parental strands, irrespective of whether the mutation has been corrected (2), or compounded (4). At middle, the tagged parents produce tagged gametes (the sex here is arbitrary). At bottom, crosses of tagged parents with members of the general population produce unilaterally tagged hybrids. Directionally biased meiotic conversion in favor of non-tagged DNA strands removes both the mutation (if present) and the tag.

When meiotic conversion is *randomly* biased, a mutation in one of the parental genomes can either be repaired or compounded (Fig 3, outcomes 1 and 3). However, this *uncertainty* in the repair process can be registered by adding an epigenetic tag symmetrically to *both* strands in the repaired region (Fig 3, outcomes 2 and 4). Subsequently, the tagged gametes can be compared with normal gametes provided by partners from the main, non-mutant, population. It would be an ‘educated bet’ (see earlier) that the conversion should result in replacement of mutant with normal sequence. So there could be a selection pressure for the conversion to be from untagged to tagged DNA strands. With PRDM9 the conversion from donor to recipient is also asymmetrical (Fig. 1). Could this experimentally observed and the hypothetical asymmetries, which both may oppose speciation, be related? Could the two asymmetries be functionally linked with retention of appropriate directionalities? Indeed, could one be the evolutionary raison d’être of the other? These questions remain with us.

### PRDM9 cannot cover all potholes

Given the great length of genome sequence in need of screening for errors, it would seem reasonable to suppose that recombination “hotspots might massively increase search efficiency by directing homology search to PRDM9 binding sites” [16]. Given this supposed advantage, why should hotspots erode? Some 25,000 to 50,000 hotspots were detected in the human genome [56]. However, only a small percentage of potential PRDM9 binding sites are used in any one meiotic cell [16, 57]. Thus, at any point in time it is not the *number* of hotspots that is limiting: “Not all PRDM9 binding sites become hotspots, and the reasons for this remain unclear” [58].

Increasing PRDM9 dosage experimentally appears to increase hotspot usage, suggesting that the availability of PRDM9 can be rate-limiting [16]. One unexplained implication of this is that an allelic pair of homologous hotspots could have PRDM9 molecules asymmetrically bound, not because of the erosion effect (Fig. 1), but because there was insufficient PRDM9. Recent work suggesting that their proper functioning requires aggregation (multimerization) of individual PRDM9 molecules, complicates this further [58].

However, another implication is that, under normal circumstances, widely dispersed, epigenetically tagged, heterologies (e.g. polymorphisms) may need to persist *for many generations* if they are to come under the surveillance of PRDM9. Failing their removal from the population by natural selection (or drift), the hotspot erosion process (Figs. 1) would serve to expose cryptic PRDM9 targets and hence, over the generations, bring different genomic segments under surveillance, so ultimately achieving repair (Fig. 2).

Then there is the pot-hole repair analogy. If a town has roads with many pot-holes and few repair vans, it makes sense to direct those vans to the most pot-holed roads. Thus, simply speeding up the pace of genome-wide surveillance might not be enough. It would seem better if close-knit collectives of epigenetically marked suspect sites could somehow (i.e. by being designated as hotspots) be recognized as needing more urgent treatment than isolated sites.

Consistent with this, Arbeithuber *et al.* note that “regions in close vicinity to these PRDM9 binding sites also showed a significant enrichment of polymorphisms in humans” [59]. While some polymorphisms are stable (presumably having escaped tagging), it seems possible that a local collective of polymorphisms that are enriched for epigenetic ‘suspect’ tags (Fig. 3), might somehow be able to attract some of the limited number of PRDM9 molecules to an otherwise cryptic hotspot in its proximity (Fig. 4).

**Figure. 4.**
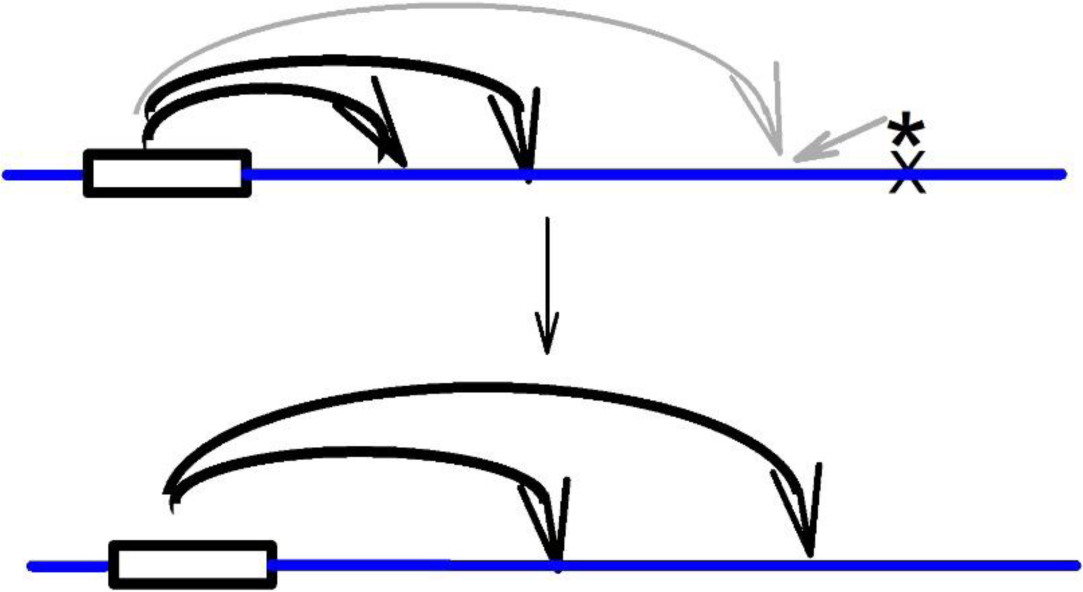
Hypothetical recruitment of a fresh PRDM9 target site by a chromosome sequence that has been epigenetically marked (asterisk) to bias meiotic conversion. Details are as in Fig. 2. Top: The curved grey arrow indicates a presumptive target site with the same sequence as those that have been targeted in previous generations (curved black arrows). The small grey arrow indicates epigenetic sequence modification prompted by some aspect of the asterisk-marked region. Bottom: The fresh target site is recognized. Given that PRDM9 concentration is limiting, one of the previous target sites is vacated. The suspect sequence with a mutation (X) has been directionally converted and the tag has been eliminated.

A hypertagged region would influence hotspot placement. Alternatively, such a collective of tagged heterologies might be able to trigger local hotspot mutation to increase affinity for PRDM9, or send a long-range signal to the *prdm9* gene that it is time to mutate its zinc fingers so as to reconfigure its global control. Whatever the ‘pothole’ search mechanism, this underlines the value of focusing recombination to hotspots, rather than randomly.

Some of the multitude of recombination-related proteins so far discovered – most not mentioned here – could be involved. Noting that increase in PRDM9 dosage only “partially rescues hybrid sterility of PWD × B6 F1 males,” Balcova *et al.* [57] distinguish the *identification* of hotspot targets from the *rate* of events subsequent to that identification, which implicate the *hstx2* gene. Their results are interpreted as “strongly indicating an independent control of global crossover rate variation and genomic crossover placement.” Given the current pace of research in this area, a role for the *hstx2* gene product could soon emerge [46].

## Conclusions

Speciation is a possible outcome of increasing variation between individuals in a species. While variation can be diminished, usually in local genome regions, by selective sweeps, there are also numerous internal mechanisms operating genome-wide to maintain DNA sequence integrity [33]. This could account for the “mysterious” selective pressure on *prdm9* [35]. The quicker *prdm9* can mutate to facilitate *symmetrical* recognition of new hotspot sequences, the quicker and more comprehensive can be the repair of heterozygosities, and the less likely the triggering of a speciation event. Since this now appears as the main role of PRDM9, then *prdm9* is best regarded as an anti-speciation gene [17, 18]. The case for this, which includes explaining the hotspot conversion paradox, is made here in the context of a chromosomal basis for hybrid sterility. To this extent, the chromosomal hypothesis is supported over genic alternatives as governing the initiation of speciation in the general case. Of course, numerous genes participate in the orchestration of meiosis and malfunction of only one can produce hybrid sterility [2]. However, in general, it is malfunction of the most distinctive feature of meiosis – the sequence-dependent pairing of homologous chromosomes – that is likely to be decisive in sparking sympatric divergence into new species.

## Acknowledgements

Virgil Reese, Root Gorelick and Rosemary Redfield gave valuable advice. Queen’s University hosts my biohistory web pages (<u>http://post.queensu.ca/~forsdyke/evolutio.htm</u> or <u>https://archive-it.org/collections/7641</u>).

The author declares no conflict of interest.

## References

1. Forsdyke DR. 2001. The Origin of Species, Revisited. McGill-Queen’s University Press, Montreal.

2. Forsdyke DR. 2010. George Romanes, William Bateson, and Darwin's "weak point." Notes Rec RS 64: 139–154.

3. Nei M, Nozawa M. 2011. Roles of mutation and selection in speciation: from Hugo de Vries to the modern genomic era. Gen Biol Evol 3: 812–829.

4. Nevo E. 2012. Speciation: chromosomal mechanisms. Encyclopedia of Life Sciences. John Wiley, Chichester, doi: 10.1002/9780470015902.a0001757.pub3

5. Kliman RM, Rogers BT, Noor MAF. 2001. Differences in (G + C) content between species: a commentary on Forsdyke’s “chromosomal viewpoint” of speciation. J Theor Biol 209: 131–140.

6. Forsdyke DR. 2004. Chromosomal speciation: a reply. J Theor Biol 230: 189–196.

7. Johannesson K. 2010. Are we analyzing speciation without prejudice? Ann N Y Acad Sci 1206: 143–149.

8. Payseur BA. 2016. Genetic links between recombination and speciation. PLoS Genet 12: e1006066.

9. Mallet J. 2006. What does Drosophila genetics tell us about speciation? Trends Ecol Evol 21: 386–393.

10. Schartl M. 2008. Evolution of Xmrk: an oncogene, but also a speciation gene? BioEssays 30: 822–832.

11. Louis EJ. 2009. Origins of reproductive isolation. Nature 457: 549–560.

12. Kao KC, Schwartz K, Sherlock G. 2010. A genome-wide analysis reveals no nuclear Dobzhansky-Muller pairs of determinants of speciation between S. cerevisiae and S. paradoxus, but suggests more complex incompatibilities. PLoS Genet 6: e1001038.

13. Hayashi K, Yoshida K, Matsui Y. 2005. A histone H3 methyltransferase controls epigenetic events required for meiotic prophase. Nature 438: 374–378.

14. Mihola O, Trachtulec Z, Vicek C, Schimenti JC, et al. 2009. A mouse speciation gene encodes a meiotic histone H3 methyltransferase. Science 323: 373–375.

15. Flachs P, Mihola O, Šimeček P, Gregorová S, et al. 2012. Interallelic and intergenic incompatibilities of the Prdm9 (Hst1) gene in mouse hybrid sterility. PLoS Genet 8: e1003044.

16. Davies B, Hatton E, Altemose N, Hussin JG, et al. 2016. Re-engineering the zinc fingers of PRDM9 reverses hybrid sterility in mice. Nature 530: 171–176.

17. Brand CA, Presgraves DC. 2016. Evolution: on the origin of symmetry, synapsis, and species. Curr Biol 26: 325–328.

18. Reese VR, Forsdyke DR. 2016. Meiotic pairing inadequacies at the levels of X chromosome, gene, or base: Epigenetic tagging for transgenerational error-correction guided by a future homologous duplex. Biol Theor 11: 150–157.

19. Ségurel L, Leffler EM, Przeworski M. 2011. The case of fickle fingers: how the PRDM9 zinc finger protein specifies meiotic recombination hotspots in humans. PLoS Biol 9: e1001211.

20. Forsdyke DR. 2017. Base composition, speciation, and why the mitochondrial barcode precisely classifies. Biol Theor 12: 157–168.

21. Forsdyke DR. 1996. Different biological species "broadcast" their DNAs at different (G+C)% "wavelengths". J Theor Biol 178: 405–417.

22. Gladyshev E, Kleckner N. 2014. Direct recognition of homology between double helices of DNA in Neurospora crassa. Nat Commun 5: 3509.

23. Gladyshev E, Kleckner N. 2016. Recombination-independent recognition of DNA homology for repeat-induced point mutation (RIP) is modulated by the underlying nucleotide sequence. PLoS Genet 12: e1006015.

24. Gladyshev E, Kleckner N. 2017. Recombination-independent recognition of DNA homology for repeat-induced point mutation. Curr Genet 63: 389–400.

25. Forsdyke DR. 2017. Speciation: Goldschmidt's chromosomal heresy, once supported by Gould and Dawkins, is again reinstated. Biol Theor 12: 4–12.

26. Danilowicz C, Lee CH, Kim K, Hatch K, et al. 2009. Single molecule detection of direct, homologous, DNA/DNA pairing. Proc Natl Acad Sci USA 106: 19824–19829.

27. Forsdyke DR. 2016. Evolutionary Bioinformatics, 3rd edn. Springer, New York.

28. Reanney DC. 1979. RNA splicing and polynucleotide evolution. Nature 277: 598–600.

29. Poole AM, Logan DT. 2005. Modern mRNA proofreading and repair: clues that the last universal common ancestor possessed an RNA genome. Mol Biol Evol 22: 1444–1455.

30. Forsdyke DR. 2007. Molecular sex: the importance of base composition rather than homology when nucleic acids hybridize. J Theor Biol 249: 325–330.

31. Montgomery TJ. 1901. A study of the chromosomes of the germ cells of metazoa. Trans Am Phil Soc 20: 154–236.

32. Winge Ö. 1917. The chromosomes, their number and general importance. Compt Rend Trav Lab Carlsberg 13: 131–275.

33. Bernstein C, Bernstein H. 1991. Aging, Sex and DNA Repair. Academic Press, San Diego.

34. Gorelick R, Heng HHQ. 2010. Sex reduces genetic variation: a multidisciplinary review. Evolution 65: 1088–1098.

35. Lesecque Y, Glémin S, Lartillot N, Mouchiroud D, et al. 2014. The red queen model of recombination hotspots evolution in the light of archaic and modern human genomes. PLoS Genet 10: e1004790.

36. Paz A, Kirzhner V, Nevo E, Korol A. 2006. Coevolution of DNA-interacting proteins and genome ‘dialect.’ Mol Biol Evol 23: 56–64.

37. Cox EC, Yanofsky C. 1969. Mutator gene studies in Escherichia coli. J Bacteriol 100: 390–397.

38. Lao PJ, Forsdyke DR. 2000. Crossover hotspot instigator (Chi) sequences in E. coli occupy distinct recombination/transcription islands. Gene 243: 47–57.

39. Forsdyke DR. 2011. The selfish gene revisited: reconciliation of Williams-Dawkins and conventional definitions. Biol Theor 5: 246–255.

40. Brick K, Smagulova F, Khil P, Camerini-Otero RD, et al. 2012. Genetic recombination is directed away from functional genomic elements in mice. Nature 485: 642–645

41. Wahls WP, Davidson MK. 2011. DNA sequence-mediated, evolutionarily rapid, redistribution of meiotic recombination hotspots. Genetics 189: 685–694.

42. Boulton A, Myers RS, Redfield RJ. 1997. The hotspot conversion paradox and the evolution of meiotic recombination. Proc Natl Acad Sci USA 94: 8058–8063.

43. Grey C, Barthès P, Chauveau-Le Friec G, Langa F, et al. 2011. Mouse PRDM9 DNA-binding specificity determines sites of histone H3 lysine 4 trimethylation for initiation of meiotic recombination. PLoS Biol 9: e1001176.

44. Bhattacharyya T, Gregorova S, Mihola O, Anger M, et al. 2013. Mechanistic basis of infertility of mouse intersubspecific hybrids. Proc Nat Acad Sci USA 110: E468–E477.

45. Bhattacharyya T, Reifova R, Gregorova S, Simecek P, et al. 2014. X chromosome control of meiotic chromosome synapsis in mouse inter-subspecific hybrids. PLoS Genet 10: e1004088.

46. Gregorova S, Gergelits V, Irena Chvatalova I, Bhattacharyya T, et al. 2017. Interference with Prdm9-controlled meiotic chromosome asynapsis overrides hybrid sterility in mice. BioRxiv preprint. http://dx.doi.org/10.1101/203505.

47. Mérot C, Salazar C, Merrill RM, Jiggins CD, et al. 2017. What shapes the continuum of reproductive isolation? Lessons from Heliconius butterflies. Proc R Soc B 284: 20170335.

48. Forsdyke DR. 2000. Haldane's rule: hybrid sterility affects the heterogametic sex first because sexual differentiation is on the path to species differentiation. J Theor Biol 204: 443–452.

49. Delph LF, Demuth JP. 2016. Haldane’s rule: genetic bases and their empirical support. J Hered 107: 383–391.

50. Oliver PL, Goodstadt L, Bayes JJ, Birtle Z, et al. 2009. Accelerated evolution of the Prdm9 speciation gene across diverse metazoan taxa. PLoS Genet 5: e1000753.

51. Harvey MG, Seeholzer GF, Smith BT, Rabosky DL, et al. 2017. Positive association between population genetic differentiation and speciation rates in new world birds. Proc Natl Acad Sci USA 114: 6328–6333.

52. Irie S, Tsujimura A, Miyagawa Y, Ueda T, et al. (2009). Single nucleotide polymorphisms in PRDM9 (MEISETZ) in patients with nonobstructive azoospermia. J Androl 30: 426–431.

53. Forsdyke DR. 2011. The B in BDM. William Bateson did not advocate a genic speciation theory. Heredity 106: 202.

54. Liénard MA, Araripe LO, Hartl DL. 2016. Neighboring genes for DNA-binding proteins rescue male sterility in Drosophila hybrids. Proc Natl Acad Sci USA 113: E4200–4207.

55. Kafer GR, Li X, Horii T, Suetake I, et al. 2016. 5-Hydroxymethylcytosine marks sites of DNA damage and promotes genome stability. Cell Rep 14: 1283–1292.

56. Myers S, Bottolo L, Freeman C, McVean G, et al. 2005. A fine scale map of recombination rates and hotspots across the human genome. Science 310: 321–324.

57. Balcova M, Faltusova B, Gergelits V, Bhattacharyya T, et al. 2016. Hybrid sterility locus on chromosome X controls meiotic recombination rate in mouse. PLoS Genet 12: e1005906.

58. Altemose N, Noor N, Bitoun E, Tumian A, et al. 2017. Human PRDM9 can bind and activate promoters, and other zinc-finger proteins associate with reduced recombination in cis. BioRxiv preprint. http://dx.doi.org/10.1101/144295.

59. Arbeithuber B, Betancourt AJ, Ebner T, Tiemann-Boege I. 2015. Crossovers are associated with mutation and biased gene conversion at recombination hotspots. Proc Natl Acad Sci USA 112: 2109–2114.

